# Basement membrane hydrogels dampen CAR-T cell activation: nanofibrillar cellulose gels as alternative to preserve T cell function in 3D cell cultures

**DOI:** 10.1101/2024.10.29.620948

**Authors:** Sonia Aristin Revilla, Alessandro Cutilli, Dedeke Rockx-Brouwer, Cynthia Lisanne Frederiks, Marc Falandt, Riccardo Levato, Onno Kranenburg, Caroline A. Lindemans, Paul James Coffer, Victor Peperzak, Enric Mocholi, Marta Cuenca

## Abstract

**Background:** Hydrogel-based 3D culture systems are emerging as a valuable tool for preclinical screening of cell-based immunotherapies against solid and hematological malignancies, such as chimeric antigen receptor T (CAR-T) cells. Hydrogels can influence T cell function in a non-desired manner due to their mechanical properties and chemical composition, potentially skewing results in preclinical testing of novel immunotherapeutic compounds.

**Methods:** In this study, we assess CD4^+^ T and CAR-T cell activation and proliferation in chemically-undefined matrices (Matrigel and basement membrane extract, BME) and compare them to a synthetic nanofibrillar cellulose (NFC) hydrogel.

**Results:** Rheometric analyses show that NFC is more rigid than Matrigel and BME. Murine CD4^+^ T cells acquire a regulatory T cell (Treg) phenotype in Matrigel and BME, while this is not observed in NFC. Proliferation and activation of human T cells are higher in NFC than in Matrigel or BME. Similarly, we show that CAR-T cell activation and proliferation is significantly impaired in Matrigel and BME, in contrast to NFC.

**Conclusions:** Our findings highlight the impact of hydrogel choice on (CAR-)T cell behavior, with direct implications for preclinical immunotherapy testing. In contrast to Matrigel and BME, NFC offers a chemically-defined 3D environment where T cell function is preserved.

**Key messages:** *What is already known on this topic:* In 3D (preclinical) tumor-killing assays for evaluating engineered T cell cytotoxicity, the surrounding matrix can influence immune cell phenotype and function, potentially skewing T cell activity. Basement membrane hydrogels such as Matrigel and basement membrane extract (BME), widely used as scaffolds for 3D culture, are inherently heterogeneous and contain extracellular matrix components that can influence lymphocyte function.

*What this study adds:* Here, we show that (CAR-)T cell function is significantly reduced in Matrigel and BME as compared to standard (2D) culture conditions. In contrast, (CAR-)T cell activity is preserved in synthetic nanofibrillar cellulose (NFC) gels. Importantly, murine T cells spontaneously acquire a Treg phenotype in Matrigel and BME. T cell proliferation and cytokine secretion are >10-fold lower in Matrigel than in NFC. Similarly, CAR-T cell survival and expansion are 10-fold higher in NFC than in Matrigel or BME.

*How this study might affect research, practice or policy:* We report that the intrinsic cytotoxic and proliferative potential of (CAR-)T cells can be underestimated when performing assays in 3D cultures based on Matrigel or BME. As an alternative, we suggest the use of chemically defined synthetic gels, and we show that nanofibrillar cellulose hydrogels are suitable 3D matrices for preserving T cell phenotype and activation.

## 1. Background

3D cell culture systems, including organoids, hold great potential for advancing biomedical developments in fundamental research, industry, and preclinical testing. These *in vitro* 3D models can be generated using healthy or diseased tissues and capture aspects of an organ’s functional and structural complexity [1]. Thus, organoids represent physiologically relevant platforms to study (cancer) cell biology and assess novel drugs, including conventional chemotherapeutics or advanced therapy medicinal products (ATMPs) such as chimeric antigen receptor (CAR)-T cells, tumor-infiltrating lymphocytes (TILs), and bispecific T cell engagers. Over the last decade, significant efforts have been made for developing 3D culture models resembling the tumor microenvironment (TME) that can be used for therapeutic screening, including cytotoxicity assays for testing novel TIL and CAR-T cell products [2–8]. The advantages of using 3D culture models for preclinical validation of tumor-targeting T cell products have been increasingly recognized in recent years. Driven by the need for therapeutic and diagnostic tools as alternatives to animal models, the global organoid market is rapidly advancing, with a predicted compound annual growth of 21.7% over the 2020-2027 period [9]. In particular, significant investments from both funding and regulatory agencies reflect a growing interest for developing robust, clinically relevant 3D culture systems for testing (engineered) T cell behavior in the context of cancer immunotherapy. Recently, the European Medicine Agency listed the implementation of organoids as tools for preclinical ATMP testing as one of the main proposals to foster ATMP development [10].

3D cultures rely on matrices made from materials such as hydrogels, polymeric scaffolds, or decellularized extracellular matrix (ECM) components to provide structural support and microenvironmental cues that shape cellular behavior and tissue development [11]. Thus, matrix choice remains a critical step for developing easy-to-implement and reliable proof-of-concept assays. Importantly, 3D culture-based preclinical drug screenings should be highly reproducible, and it is therefore relevant to base such models on matrices with a defined composition that do not interfere with cell phenotype and activation in an uncontrolled way. Currently, 3D culture models based on animal-derived ECM protein mixtures such as Matrigel or basement membrane extract (BME) are widely used [12,13]. Matrigel and BME are obtained from a murine tumor (EHS sarcoma) which must be propagated in mice; at least 25 tumor-bearing mice are needed to prepare a single liter of matrix [14]. Besides sustainability concerns, a significant limitation of Matrigel and BME is their undefined and variable composition, with unknown amounts of ECM proteins and growth factors [12,13].

In the context of 3D (preclinical) tumor-killing assays for evaluating engineered T cell cytotoxicity, the surrounding matrix can influence immune cell phenotype and function, potentially skewing T cell activity. Animal-derived matrices such as Matrigel and BME are inherently heterogeneous and contain diverse ECM components that can be present at variable concentrations. These factors include insulin growth factor-1 (IGF-1), which stimulates T cell growth and differentiation [15]; transforming growth factor-β (TGF-β), which drives the proliferation, differentiation, and expansion of regulatory T (Treg) cells fostering their development toward an immunosuppressive phenotype [16]; and vascular endothelial growth factor (VEGF), which can inhibit T cell antitumor activity and promote Treg cells differentiation [16]. In addition, it was recently reported that ECM viscoelasticity regulates T cell phenotype – imprinting long-term features – and function, since tuning ECM viscoelasticity results in the generation of functionally distinct T cells [17]. Thus, to assess T cell function in 3D culture models, it is necessary to use matrices with defined viscoelasticity and devoid of uncontrolled factors that can skew T cell activation and proliferation. This holds particular significance in preclinical ATMP validation, where the ability of a candidate T cell therapeutic product to proliferate and demonstrate anti-tumor activity is critical in determining whether to advance or discontinue further clinical development.

In light of these considerations, the development of next-generation 3D culture systems for assessing T cell function requires a chemically defined hydrogel platform, exempt from undefined components and sourced from non-animal origins. Additional properties that would be desirable in this matrix include commercial availability, ease of implementation and handling (for instance with the possibility to embed cells at room temperature), and compatibility with downstream cellular analysis by flow cytometry, PCR, or RNA sequencing. In this study we analyze a nanofibrillar cellulose (NFC) hydrogel as a chemically defined alternative to Matrigel or BME with all the abovementioned features. The nanofibrillar cellulose fibers self-assemble forming a random 3D structure to support cell growth, differentiation, and spheroid formation [18,19]. Here, we show that culture of murine T cells in Matrigel and BME leads to an increase in the proportion of Treg cells, while this is not observed in NFC. In addition, human CD4^+^ T cells and CAR-T cells exhibit significantly higher activation and proliferation in NFC than in animal-derived matrices, although short-term CAR-T cell cytotoxicity was preserved in all hydrogels. In conclusion, NFC gels provide a chemically-defined alternative to Matrigel and BME to preserve (CAR-)T cell effector functions in 3D culture models.

## 2. Results

### 2.1. NFC hydrogel is stiffer than Matrigel and BME

First, the viscoelastic behavior of NFC hydrogels, Matrigel and BME was assessed using oscillatory rheological analysis. A time/temperature ramp was used as to analyze the crosslinking kinetics and stiffness of the viscoelastic hydrogel materials, since both Matrigel and BME display a temperature-dependent gelation mechanism (**Figure 1a**). Matrigel and BME displayed similar crosslinking kinetics. The results also showed a higher stiffness of NFC hydrogel when compared to Matrigel or BME. Moreover, the storage modulus of NFC is invariant with the temperature, indicating that the material is already a stable gel at room temperature. These findings have direct implications in the handling of the materials for cell culture assays, as Matrigel and BME need to be manipulated at cold temperatures when encapsulating cells not to trigger unwanted thermal crosslinking too early, whereas for NFC cells can be mixed in the gel at any temperature relevant for cell viability.

**Figure 1.**
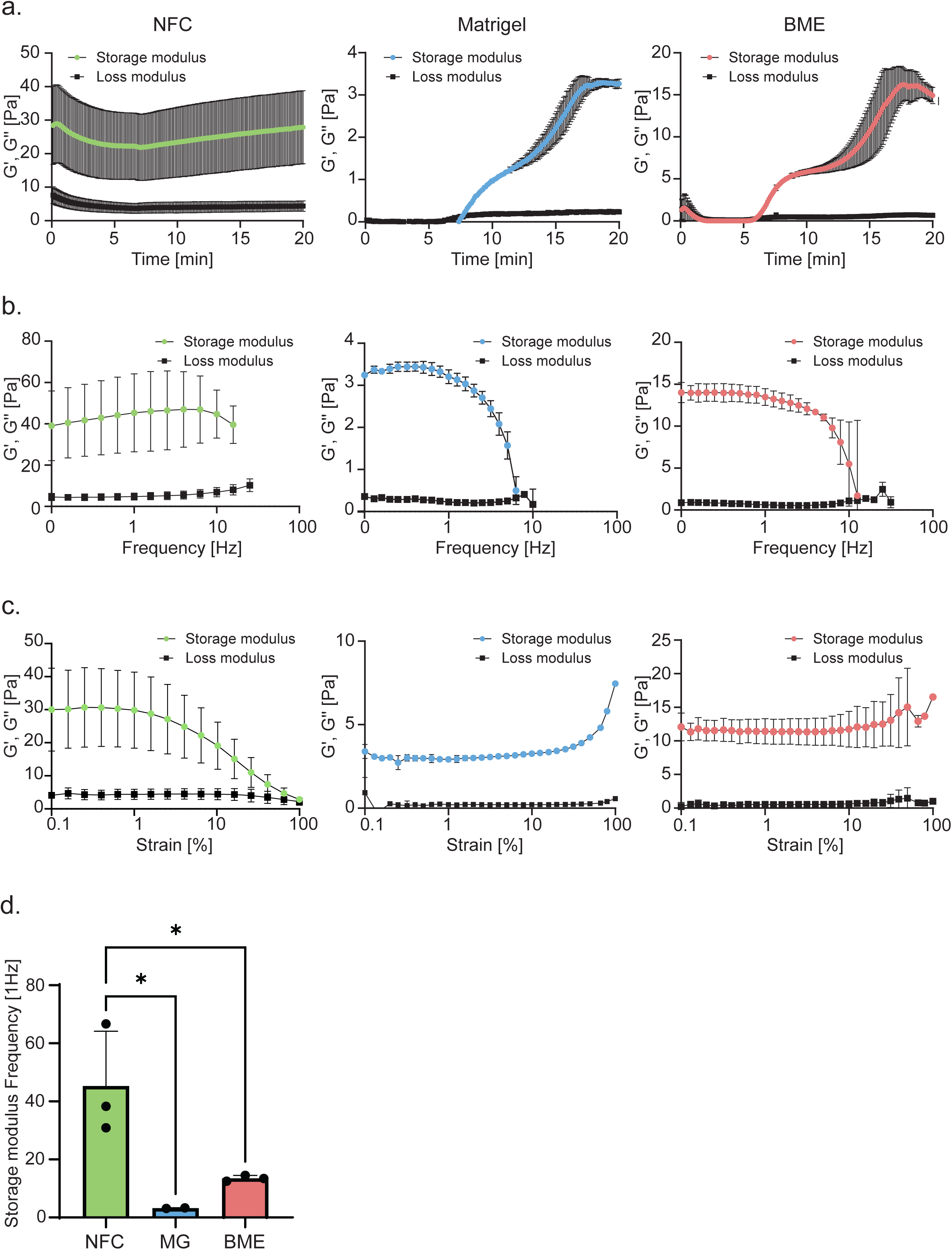
Rheological properties of NFC hydrogel, Matrigel and BME. (**a**) Rheological time and temperature sweep measurements displaying the crosslinking kinetics and behavior of NFC, Matrigel (MG) and BME. (**b**) Rheological frequency sweep measurements at 37°C, showing the frequency dependent viscoelastic behavior of NFC, Matrigel and BME. (**c**) Rheological amplitude sweep at 37°C, showing the amplitude dependent viscoelastic behavior of NFC, Matrigel and BME. Data representative of three independent measurements are presented as means ± SD. Data were analyzed using unpaired one-way ANOVA with Tukey’s multiple comparison test. (*P < 0.05). NFC, nanofibrillar cellulose hydrogel; MG, Matrigel; BME, basement membrane extract.

Next, a frequency sweep was applied to the materials to study the time dependent behavior of the hydrogels in response to a non-destructive deformation (**Figure 1b**). These results showed that especially NFC seems to be more resilient to faster deformations than Matrigel and BME, while Matrigel showed the least resilience to fast deformations. To further characterize the hydrogels, an amplitude sweep was performed to assess the range of strain the materials can withstand before yielding (**Figure 1c**). Both Matrigel and BME showed a strain stiffening behavior when subjected to shear strain values >50%. Conversely, in line with the particulate nature of the hydrogel, NFC displayed a reduction in storage modulus with increasing shear amplitude, suggesting that elevated strains promote a more fluid-like behavior of the gel by disrupting the interactions that keep the NFC slurry cohesive. This aspect can be beneficial to mechanically disaggregate the gel to retrieve embedded cells. Finally, to provide a comparison of the stiffness values of the different hydrogels, we extracted the storage moduli of the materials at 1 Hz (**Figure 1d**). The values derived from the frequency sweep assays demonstrate that NFC is significantly stiffer than both Matrigel and BME, although all gels present values comparable to soft tissues, in the range between 3-40 Pa of shear storage moduli, typically suitable for cell culture and to perform migration and invasion assays.

### 2.2. Culture of murine CD4^+^ T cells in Matrigel and BME leads to an increase in Treg cell numbers, while this is not observed in NFC hydrogels

We next explored how NFC, Matrigel and BME impact the activation of embedded murine CD4^+^ T cells isolated from transgenic C57BL/6 Foxp3eGFP mice. In this mouse model, eGFP expression is controlled by the promoter of *Foxp3*, the key transcription factor driving Treg cell differentiation and function [20,21]. Consequently, eGFP serves as a surrogate marker for the identification of Treg cells. T cell activation involves a three-step process: first, antigen recognition through the TCR; second, CD28 co-stimulation; and finally, cytokine-mediated triggering of interleukin-2 receptor (IL-2R or CD25) signaling. In our experimental set-up, T cells were stimulated *in vitro* using anti-CD3/CD28 monoclonal antibodies along with soluble IL-2. T cell activation induces changes in their morphology with an increase in size and formation of clusters, which are visible under the microscope. Notably, after five days in culture, CD4^+^ T cells cultured in 2D and in NFC exhibited these characteristic clusters, while this was not observed for CD4^+^ T cells embedded in Matrigel and BME (**Figure 2a**). Next, CD4^+^ T cell viability in the different gels was evaluated by flow cytometry. We found that all conditions yielded a similar number of viable cells (**Figure 2b**). We then examined additional parameters related to CD4^+^ T cell activation such as CD25 expression, which is upregulated in activated T cells. We observed that CD25 expression was significantly reduced when CD4^+^ T cells were cultured in BME (MFI 370 ± 24) as compared to NFC (560 ± 125) or Matrigel (590 ± 207), suggesting reduced T cell activation (**Figure 2c**).

**Figure 2.**
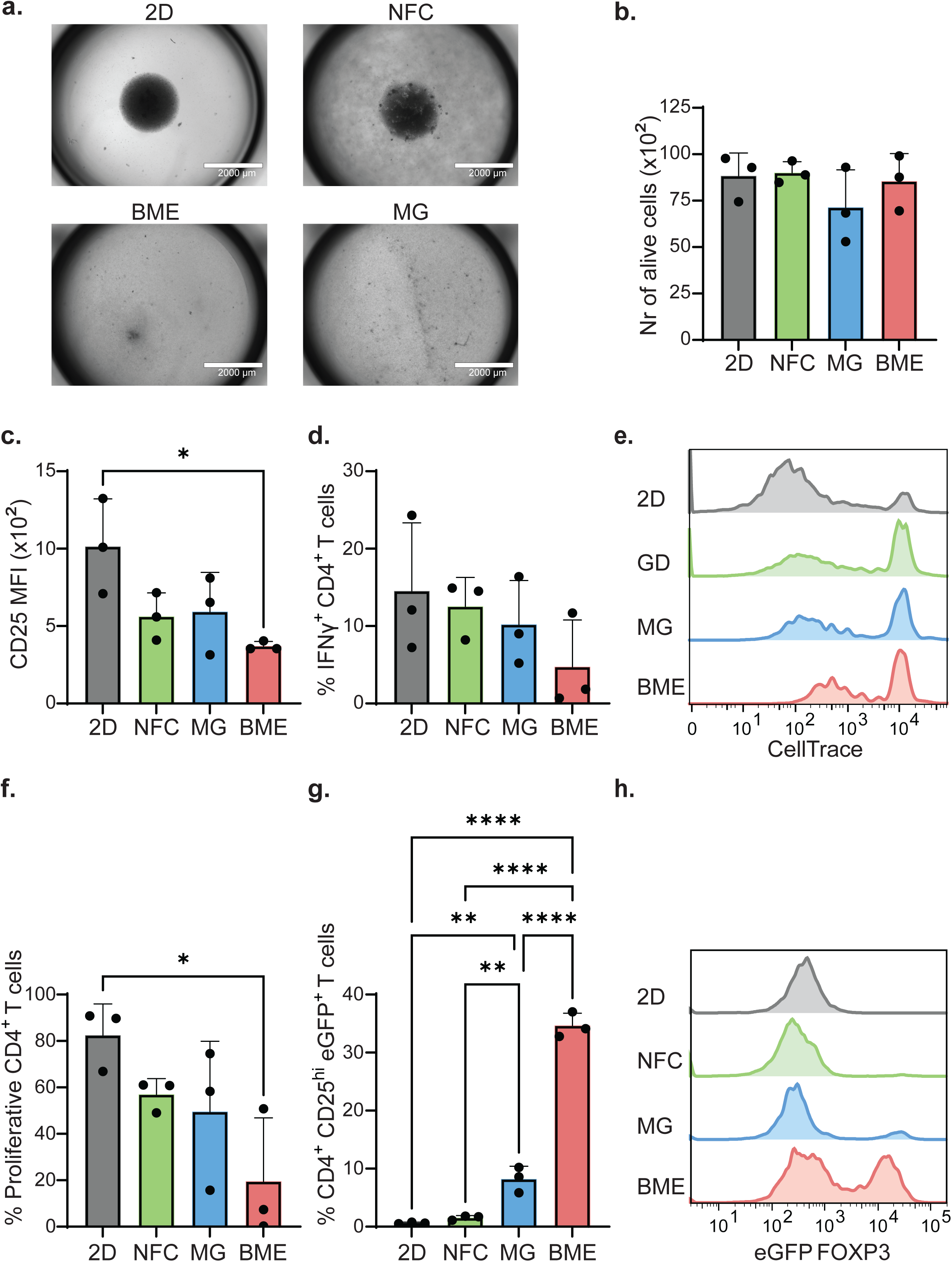
ECM-derived hydrogels (Matrigel and BME) influence murine CD4^+^ T cell activation, function and differentiation to regulatory T cells. (**a**) Representative bright field images (EVOS microscope, scale bar 2000 uM) of CD4^+^ T cells purified from Foxp3eGFP mice stimulated ex-*vivo* with anti-CD3 (1 μg/ml) and anti-CD28 (1 μg/ml) in standard 2D suspension (control) or embedded in different hydrogels (NFC, Matrigel and BME). CD4^+^ T cells were analyzed by flow cytometry after 5 days in culture. (**b**) Number of viable CD4^+^ T cells recovered after culture. (**c**) Median fluorescent intensity (MFI) of CD25 in CD4^+^ T cells. (**d**) Percentage of IFNγ-expressing CD4^+^ T cells. (**e**) Representative histograms showing CellTrace Violet expression on CD4^+^ T cells and (**f**) proportion of proliferative CD4^+^ T cells (CTV-low). (**g**) Percentage of Foxp3 eGFP^+^ CD25^hi^ Treg cells and (**h**) representative histograms showing Foxp3 expression on Foxp3-eGFP^+^ Treg cells. Each point represents an individual mouse. Data from three independent experiments are presented as means ± SD. Data were analyzed using unpaired one-way ANOVA with Tukey’s multiple comparison test. (*P < 0.05, **P < 0.005, ****P < 0.0001.). NFC, nanofibrillar cellulose; MG, Matrigel; BME, basement membrane extract.

To further investigate functional aspects of T cell activation, we assessed the fraction of CD4^+^ T cells expressing interferon gamma (IFNγ) using flow cytometry. Our analysis showed similar proportions of IFNγ^+^ CD4^+^ T cells among the different culture conditions (**Figure 2d**), suggesting that the functional capacity of CD4^+^ T cells to produce this cytokine is preserved independently of the matrix used, although there was a non-significant trend towards decreased IFNγ production in BME. In parallel, CD4^+^ T cell proliferation was evaluated by measuring CellTrace Violet dilution by flow cytometry. Proliferation of CD4^+^ T cells embedded in BME was significantly decreased (19.5% ± 22% proliferative cells) compared to proliferation in 2D (82.5% ± 11%), NFC (57% ± 6%) and Matrigel (49.5% ± 25%) (**Figure 2e-f**). Last, we analyzed the proportion of CD4^+^ T cells expressing Foxp3 eGFP within our cultures to assess changes in Treg numbers in the different matrices. A significant increase in the percentage of Treg cells (defined as Foxp3 eGFP^+^ CD25hi CD4^+^ T cells) was observed when activated CD4^+^ T cells were embedded in Matrigel (8.2% ± 1.8%) or BME (34.6% ± 1.8%) for five days. In contrast, embedding activated CD4^+^ T cells in NFC did not increase Treg cell numbers (**Figure 2g-h and Figure S1**). Taken together, these data highlight that ECM-derived matrices such as Matrigel and BME can affect murine CD4^+^ T cell activation, proliferation and differentiation towards a regulatory phenotype, while this is not observed in NFC gels.

### 2.3. Increased human CD4^+^ T cell viability, activation, and proliferation in NFC compared to ECM-derived hydrogels

Next, we explored how NFC, Matrigel and BME influence the phenotype and function of human CD4^+^ T cells. We examined viability, activation markers, proliferation, and spontaneous Treg induction in the different hydrogels after five days in culture. CD4^+^ T cells were isolated from cord blood of human donors and stimulated *in vitro* using anti-CD3/CD28 monoclonal antibodies along with soluble IL-2, following the same protocol as for murine T cells. CD4^+^ T cell viability was significantly higher in NFC than in Matrigel and BME (**Figure 3a**). T cell activation, as measured by CD25 expression, was lower in Matrigel (MFI 87 ± 7.7) and BME (107 ± 16) than in NFC (225 ± 117) and 2D controls (363 ± 93) (**Figure 3b-c**). To functionally evaluate activated CD4^+^ T cells in the different matrices, the percentage of CD4^+^ T cells expressing IFNγ was measured by flow cytometry. CD4^+^ T cells cultured in NFC produced more IFNγ (28.3% ± 16% IFNγ^+^ cells) than cells cultured in Matrigel (2.5% ± 2.6%) and BME (0.9% ± 0.1%) (**Figure 3d-e and Figure S2**). In addition, CD4^+^ T cell proliferation in NFC was also significantly higher (39% ± 21% proliferative cells) than in Matrigel (4.7% ± 4.5%) or BME (1.6% ± 0.19%) (**Figure 3f-g**). In contrast to murine CD4^+^ T cells, we found that neither Matrigel nor BME promoted an increase in Treg cell numbers (**Figure 3h**). Taken together, our findings demonstrate that survival and activation of human CD4^+^ T cells is higher in synthetic NFC gels than in ECM-derived matrices.

**Figure 3.**
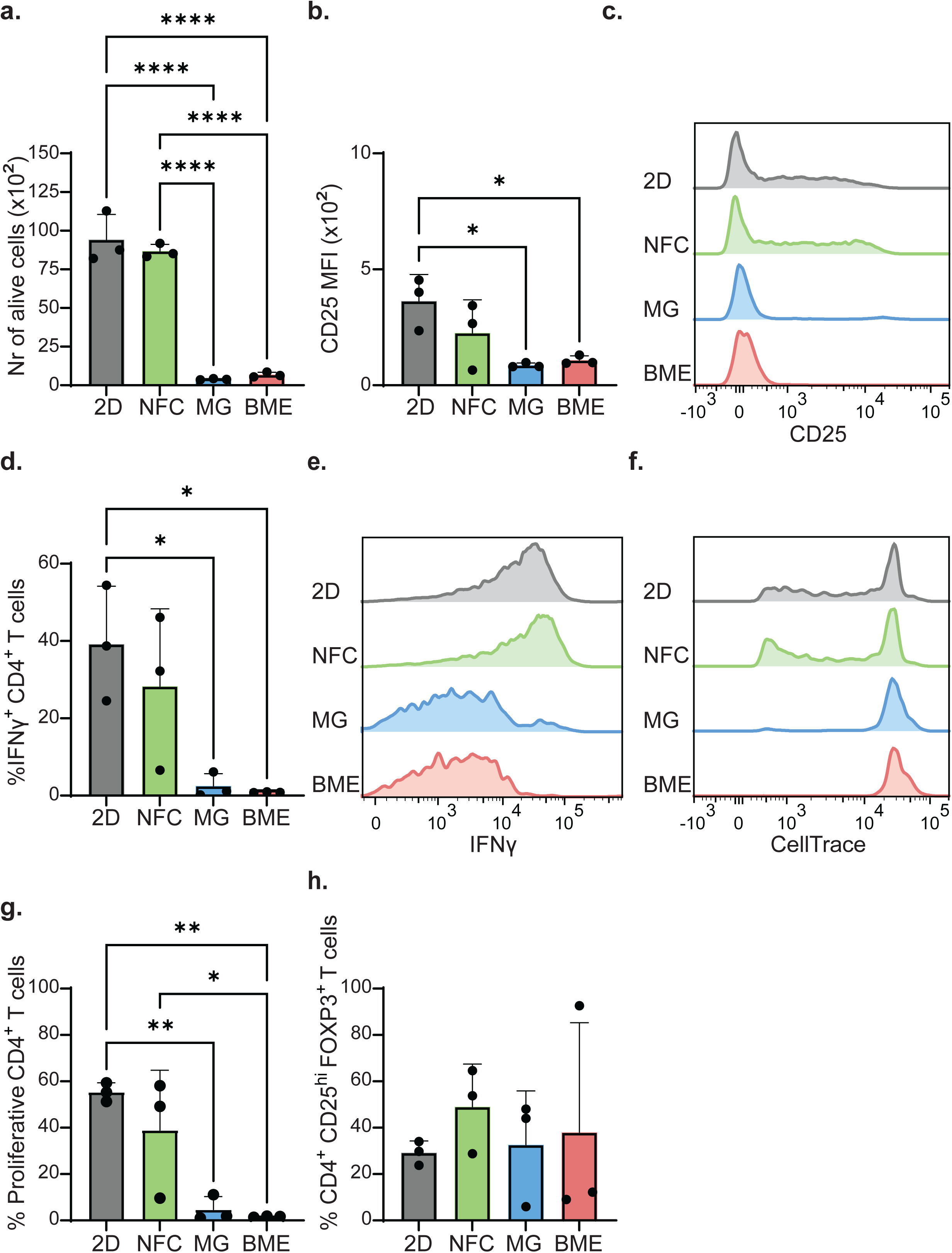
ECM-derived hydrogels (Matrigel and BME) hamper human CD4^+^ T cell activation and function. (**a-h**) CD4^+^ T cells isolated from cord blood mononuclear cells were stimulated *ex-vivo* with anti-CD3 (1 μg/ml) and anti-CD28 (1 μg/ml) for 5 days in standard 2D suspension (control) or embedded in NFC, Matrigel and BME. (**a**) Number of viable CD4^+^ T cells recovered after culture. (**b**) Median fluorescent intensity (MFI) of CD25 in CD4^+^ T cells and (**c**) representative histograms showing CD25 expression on CD4^+^ T cells. (**d**) Percentage of IFNγ-expressing CD4^+^ T cells and (**e**) representative histograms showing IFNγ expression on CD4^+^ T cells. (**f**) Representative histograms showing CellTrace Violet expression on CD4^+^ T cells and (**g**) proportion of proliferative CD4^+^ T cells (CTV-low). (**h**) Percentage of FOXP3^+^ CD25^hi^ Treg cells. Each point represents an individual donor. Data from three independent experiments are presented as means ± SD. Data were analyzed using unpaired one-way ANOVA with Tukey’s multiple comparison test. (*P < 0.05, **P < 0.005, ****P < 0.0001). NFC, nanofibrillar cellulose; MG, Matrigel; BME, basement membrane extract.

### 2.4. CAR-T cell cytotoxicity in short term cultures is comparable between NFC, Matrigel and BME

CAR-T cells are engineered immune cells designed to identify and eradicate cancer cells by recognizing distinct tumor-associated antigens. *In vitro* assays testing CAR-T cell cytotoxicity against cancer cells are critical for understanding the biology of the interactions between CAR-T and tumor cells, and for screening new drug candidates. Here, we evaluated how NFC, Matrigel and BME impact the cytotoxicity of CD20 CAR-T cells against Daudi cells, a Burkitt’s lymphoma cell line. In our experiments, CD20 CAR-T cells were co-cultured with Daudi cells at various effector-to-target ratios (E:T) in each hydrogel. After 24 hours of co-culture, the viability of CD20 CAR-T and Daudi cells was analyzed by flow cytometry. A comparable number of viable CD20 CAR-T cells was recovered across the different culture conditions (**Figure 4a-b**). Regarding CAR-T cytotoxicity, a dose-dependent correlation between increasing E:T ratios and target cell apoptosis could be observed in all matrices. CAR-T mediated killing was comparable in the three hydrogels (**Figure 4c**). Overall, these findings indicate that CAR-T mediated cytotoxicity in short-term (24h) co-cultures is preserved in NFC, Matrigel and BME.

**Figure 4.**
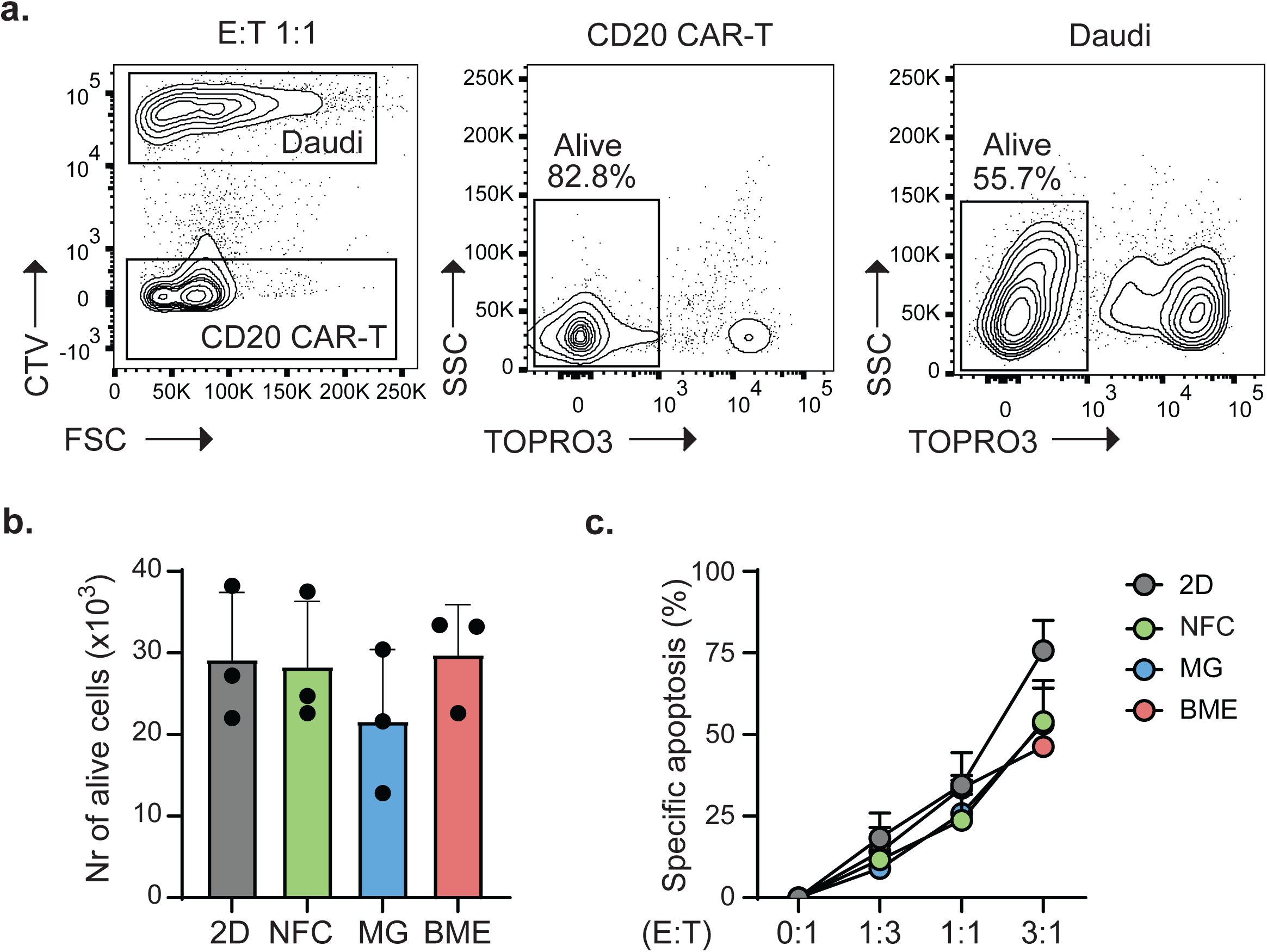
CD20 CAR-T cell cytotoxicity in NFC, Matrigel and BME. CD20 CAR-T cells were co-cultured with Daudi (Burkitt lymphoma cell line) cells labelled with CellTrace Violet (CTV) for 24h at the different effector to target (E:T) ratios specified, in standard 2D suspension (control) or embedded in the specified hydrogels. (**a**) Representative gating strategy for analyzing the viability of Daudi and CD20 CAR-T cells by flow cytometry. (**b**) Number of viable CAR-T cells recovered after culture in the 1:1 E:T condition (50.000 Daudi + 50.000 CAR-T cells/well) as measured by flow cytometry using counting beads (Flow-Count fluorospheres). Data were analyzed using a one-way ANOVA with Tukey’s multiple comparison test. (**c**) Specific lysis (%) of Daudi cells induced by CD20 CAR-T cells at the specified E:T ratios. Specific apoptosis was calculated by applying the following formula: [(%viable untreated - %viable treated)/ %viable untreated] x 100. Data from three independent experiments are presented as means ± SD. Data were analyzed using a two-way ANOVA with Tukey’s multiple comparison test. NFC, nanofibrillar cellulose; MG, Matrigel; BME, basement membrane extract.

### 2.5. CAR-T cell activation and proliferation is preserved after a 4-day stimulation in NFC, but not in Matrigel or BME

Since preclinical CAR-T cell testing often involves culture periods longer than 24h, we evaluated how the activation and functionality of CD20 CAR-T cells evolves over longer-term cultures in different 3D matrices. To address this, we cultured CD20 CAR-T cells in NFC, Matrigel and BME for 4 days in the presence of IL-2, anti-CD3 and anti-CD28 monoclonal antibodies to induce proliferation. Morphological changes associated with cell activation, such as cell enlargement and the formation of distinctive clusters, were particularly pronounced in CAR-T cells cultured in NFC (**Figure 5a**). The number of alive CD20 CAR-T cells retrieved from the different culture conditions was quantified by flow cytometry. CAR-T cell viability was significantly reduced in BME (27,706 ± 11,315 alive cells) as compared to NFC (72,836 ± 18,082), 2D or Matrigel (**Figure 5b**). In line with our previous findings, CD25 expression was significantly lower when CAR-T cells were cultured in Matrigel (MFI 500 ± 401) and BME (407 ± 251) compared to NFC (5,194 ± 1,528) (**Figure 5c-d**). In addition, CAR-T cells embedded in NFC displayed higher proliferation rates (46.8% ± 7.2% proliferative cells) than in Matrigel (4.5 ± 5.1) or BME (0.9 ± 2.6) (**Figure 5e-f**). Thus, our data indicates that over longer culture periods, ECM-derived hydrogels can negatively impact CAR-T cell activation and proliferation.

**Figure 5.**
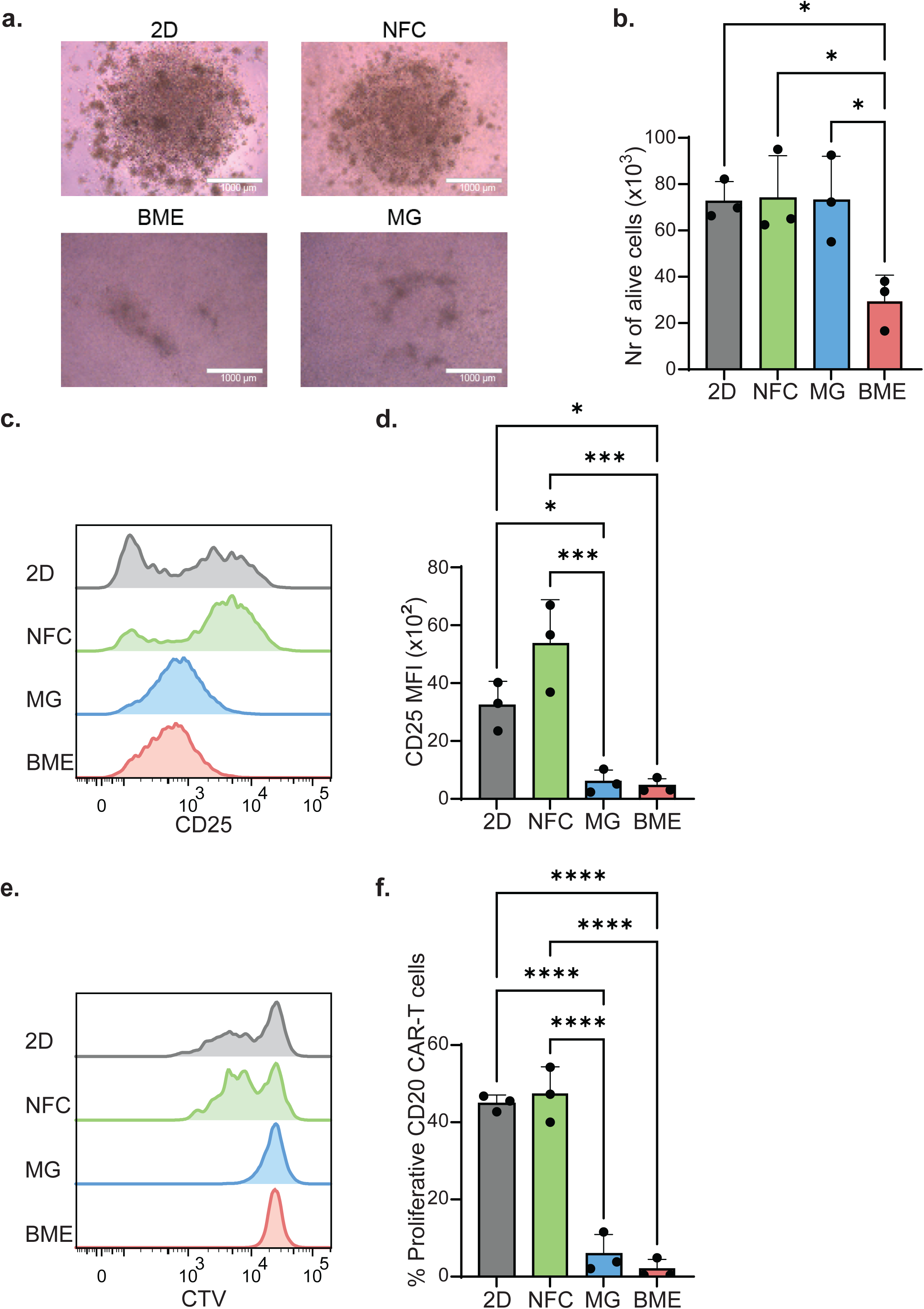
ECM hydrogels (Matrigel and BME) reduce CAR-T cell activation and proliferation. (**a**) Representative bright field images (scale bar, 1000 μM) of CD20 CAR-T cells stimulated *in vitro* for 4 days with coated anti-CD3 (1.6 μg/ml) and soluble anti-CD28 (1 μg/ml) in either standard 2D suspension (control) or the indicated gels. (**b**) Number of viable CAR-T cells recovered after culture as measured by flow cytometry using counting beads (Flow-Count fluorospheres). (**c**) Representative histograms showing CD25 expression in CAR-T cells and (d) median fluorescence intensity (MFI) of CD25 in CAR-T cells. (**e**) For flow cytometry-based analysis of proliferation, CD20 CAR-T cells were labelled with CellTrace Violet (CTV) dye on day 0 and stimulated as indicated in (**a**). Representative histograms showing CTV expression on CAR-T cells and (**f**) proportion of proliferative CAR-T cells (low CTV expression). Data from three independent experiments are presented as means ± SD. Data were analyzed using unpaired one-way ANOVA with Tukey’s multiple comparison test. (*P < 0.05, ***P < 0.0005, ****P < 0.0001). NFC, nanofibrillar cellulose; MG, Matrigel; BME, basement membrane extract.

## 3. Discussion

3D culture systems are more versatile than 2D monolayer cultures, offering the possibility to mimic more accurately different cellular environments [17,22–25]. Still, establishing 3D culture platforms that are at the same time reproducible, high-throughput and compatible with both tumor and immune cell functions remains a challenge for the development of (personalized) models for testing immunotherapeutic drugs. It is important to ensure that the chosen matrix does not interfere with immune cell function in an uncontrolled manner. In this study, we compare (CAR-)T cell activation and proliferation in Matrigel, BME, and NFC, a nanocellulose-based synthetic hydrogel. Despite the effectiveness of Matrigel and BME in culturing several types of organoids, animal-derived matrices are limited by their non-defined composition, with varying amounts of ECM components and growth factors leading to reproducibility issues [12,13,26,27]. Here, we explore NFC as a chemically-defined alternative and propose it as a solution to prevent uncontrolled skewing of T cell phenotype in 3D cultures. T cell phenotype *in vivo* is influenced by the anatomical location, motivating recent studies to optimize culture conditions and matrix composition for investigating T cell biology *in vitro* [17,22,24,25,28,29]. Previous reports have explored the suitability of chemically and non-chemically defined matrices as 3D scaffolds for culturing T cells, showing superior T cell expansion in 3D as compared to standard 2D culture [25]. Importantly, the mechanical properties of hydrogels, such as stiffness and viscoelasticity, affect T cell phenotype and activation [17,24,25,30,31]. Matrix stiffness modulates TCR mechanotransduction, regulating T cell activation and proliferation [24,25,30,31]. Stiffer 3D environments generally lead to increased CD4^+^ T cell activation and expansion [25,30,31]. Additionally, matrix viscoelasticity can induce transcriptome changes, such as CD25 upregulation, directing the generation of functionally distinct T cell populations [17]. In line with previous findings, we also observed that both matrix stiffness and composition influence murine and human CD4^+^ T cell behavior. Gel rheology analyses revealed that NFC is significantly stiffer than Matrigel or BME (**Figure 1**). Importantly, we report for the first-time induction of murine Treg cells when CD4^+^ T cells were embedded within Matrigel and BME, while this was not observed in NFC (**Figure 2g-h**). This finding aligns with a previous study where murine CD4^+^ T cell activation was analyzed in PDMS substrates with variable stiffness, showing a significant increase in Treg cell numbers on lower stiffness substrates [32]. In addition, our data showed enhanced viability, activation and expansion of human CD4^+^ T cells embedded in NFC compared to Matrigel and BME (**Figure 3a-g**), supporting the notion that stiffer hydrogels favor T cell activation. In contrast to murine T cells, we did not observe Treg induction from naïve human CD4^+^ T cells in Matrigel or BME (**Figure 3h**). This discrepancy could be explained by species-specific responses to matrix composition. Matrigel and BME are basement membrane extracts obtained from a mouse sarcoma, and therefore it is possible that murine CD4^+^ T cells are more sensitive to the biochemical cues present in these gels than human T cells. Matrigel and BME contain transforming growth factor (TGF)-β and vascular endothelial growth factor (VEGF), which can induce murine Treg cells [13,16,26]. Moreover, both TGF-β and VEGF can inhibit CD4^+^ T cell activation [13,16,26], which could underlie the observed absence of activation in human CD4^+^ T cells within Matrigel and BME matrices. Taken together, our data highlights the need for considering both matrix composition and stiffness when establishing 3D cell culture systems for analyzing T cell biology.

This is the first study addressing how CAR-T cell performance can be influenced by the 3D culture matrix. (Immuno)therapy testing in high-throughput, reproducible 3D models for solid and hematologic malignancies is increasingly becoming an essential step in preclinical development [33,34]. Matrigel and BME are widely used as hydrogel scaffold for organoid research [2,35], including 3D models for assessing CAR-T or TIL cytotoxicity [36–40]. Here, we show that CAR-T cell activation and proliferation after anti-CD3 stimulation can be severely impaired in Matrigel and BME, and this reduction was not observed when CAR-T cells were cultured in NFC (**Figure 5**). CAR-T cell cytotoxicity was comparable across all hydrogels (**Figure 4**), suggesting that, while Matrigel and BME may hinder the downstream TCR signaling events leading to T cell expansion, CAR-mediated recognition is preserved. In the context of tumor immunology, previous reports have shown upregulation of the inhibitory receptors PD-1 and CTLA-4 in CD4^+^ TILs cultured with Matrigel-inlaid melanoma organoids [37]. In an *in vivo* mouse model, co-implantation of the breast cancer cell line 4T1 with Matrigel, as opposed to implantation in PBS, was reported to impair T cell migration, activation, and function within the xenograft tumors [41]. Matrigel reduced *in vivo* T cell infiltration in the tumors, decreased the percentage of IFNγ secreting T cells, and abolished T cell activation as measured by CD69 and CD44 expression [41]. This study showed that laminin-111, one of the most abundant matrisome proteins in Matrigel [42], significantly inhibits T cell activation and proliferation [41]. Of note, in most mature tissues, laminin-111 is not as abundant as in Matrigel [43]. Matrigel-induced immunosuppression is also driven by the presence of collagen. Collagen type IV represents ∼30% of the total Matrigel composition [44]. Multiple collagen types, including collagen type I, IV, and VI, can drive T cell exhaustion by binding the leukocyte-specific collagen receptor LAIR1 and impairing T cell activation [45]. Thus, CAR-T cell activation in Matrigel can be skewed by the presence of different combinations of collagen subtypes in variable amounts. Of note, the ECM is dynamic and tissues have unique ECM compositions adapted to their needs, which can be further modified in pathological conditions[46]. Collagen and laminin gene expression profiles are highly tissue- and tumor-specific [42,47]. Therefore, despite being rich in collagen and laminin, the mouse sarcoma-derived ECM present in Matrigel and BME does not provide a representative niche for either tumors or healthy tissues. In contrast, NFC gels can be supplemented on demand with relevant ECM components to recreate tissue-specific matrices [48]. Importantly, CAR-T cell performance is also influenced by their intrinsic phenotype and differentiation stage. Successful clinical response to CAR-T cell therapy has been associated to CAR-T upregulation of genes related to a memory cell phenotype [49]. Previous reports show that after a 5-day culture, T cells embedded in Matrigel express significantly lower levels of the memory marker CD45RO than cells cultured in 2D or a 3D polystyrene scaffold [25]. Together with our observations of reduced CAR-T cell proliferation and CD25 expression in Matrigel and BME as compared to NFC, these data highlight the relevance of using chemically defined hydrogels for testing T cell tumor reactivity.

In summary, we report that the intrinsic cytotoxic and proliferative potential of (CAR-)T cells can be underestimated when performing assays in 3D cultures based on Matrigel or BME. As an alternative, we suggest the use of chemically defined synthetic gels, and we show that nanofibrillar cellulose hydrogels are suitable 3D matrices for preserving T cell phenotype and activation.

## 4. Methods

### 4.1. Time/Temperature sweep, Frequency sweep, Amplitude sweep

Rheology experiments on gel precursor solutions to determine the crosslinking kinetics were assessed using a DHR2 rheometer (TA Instruments, The Netherlands). For rheology assays, gel precursors were prepared at the same concentrations used for T cell culture as described below (Matrigel and BME were diluted 1:1 in cell culture medium, and 1.5% nanofibrillar cellulose hydrogel (GrowDex) was diluted to 0.25% w/v in culture medium). Time/temperature sweep experiments were performed at a frequency of 1.0 Hz, angular frequency of 6.283 rad/s, with 5.0% constant strain starting at 4°C with an increase of 5°C per minute until 37°C was reached, then the temperature kept constant for the duration of the experiment (n = 3 independent samples). Subsequently, frequency sweep experiments were performed at a frequency range from 0.1 Hz to 100 Hz with 5.0% constant strain at 37°C. Subsequently, amplitude sweep experiments were performed at a frequency of 1.0 Hz at a strain rate from 0.1% to 100% strain at 37°C. A volume of 100 µL of gel was used with a gap size of 300 µm. A 20.0 mm parallel EHP stainless steel plate was used as geometry.

### 4.2. Murine CD4^+^ T cell isolation

CD4^+^ T cells were obtained from transgenic B6-Foxp3^EGFP^ mice (B6.Cg-*Foxp3^tm2(EGFP)Tch^*^/J^, The Jackson Laboratory, stock no. 006772). Lymph nodes (inguinal, brachial, axillary, and cervical) and spleens were harvested and smashed against a cell strainer to obtain single-cell suspensions. Cells were centrifuged at 400g for 4 minutes at 4°C and resuspended in MACS buffer (2% heat-inactivated FBS (Gemini Bio-Products) and 2mM EDTA in PBS), followed by cell counting using a TC20 automated cell counter (Bio-Rad). CD4^+^ T cells were isolated using mouse CD4 (L3T4) microbeads (Miltenyi Biotec) according to the manufacturer’s protocol. Each mouse sample was processed with one LC column (Miltenyi Biotec) tightly placed in the QuadroMACS™ Separator (Miltenyi Biotec).

### 4.3. Human CD4^+^ T cell isolation

CD4^+^ T cells were isolated from cord blood of healthy donors obtained according to the Declaration of Helsinki. The collection protocol was approved by the Ethical Committee of the Utrecht Medical Center (UMCU) in the Netherlands. After Ficoll-Paque (GE Healthcare) gradient separation, cord blood mononuclear cells (CBMCs) were cryopreserved for later use. CD4^+^ T cells were isolated from the CBMCs fraction in MACS buffer using the MagniSort human CD4^+^ T cell enrichment kit (Thermo Fisher) and BD IMag Cell Separation Magnet (BD Biosciences) following the manufacturer’s protocol.

### 4.4. Cell encapsulation in NFC, Matrigel and BME

Cell suspensions were spun down (1500 rpm, 5 minutes, 4 °C) and pellets were resuspended in either NFC (GrowDex, UPM Biomedicals), Matrigel (Corning) or BME (Cultrex® RGF BME Type 2, R&D systems). GrowDex (supplied as a suspension of 1.5% nanofibrillar cellulose in 98.5% ultra-pure water) was diluted to 0.25% in culture medium, using low-retention pipet tips. Cell pellets were resuspended in 70 μL of this 0.25% NFC suspension, and distributed in 96-well-U plates. 130 μL of culture medium supplemented with soluble IL-2 (20 U/ml, PeproTech) were carefully pipetted on top of the gel. For encapsulation in Matrigel and BME, micro centrifuge tubes containing the cell pellets were placed on ice and resuspended in 70 µL of cold Matrigel or BME diluted 1:1 in culture medium. Gel crosslinking was induced by incubating the gel precursors at 37 °C for 30 min. After this time, 130 µL of medium supplemented with soluble IL-2 (20 U/ml, PeproTech) was carefully pipetted on top of the gels. After culture, cells were recovered from NFC hydrogels by adding GrowDase (UPM Biomedicals) (800 μg GrowDase per mg of GrowDex) and incubating at 37 °C for 3h. Matrigel and BME were disrupted by using Cell Recovery Solution (Corning). All samples were filtered through a 30 μM cell strainer before flow cytometric analysis.

### 4.5. Culture of murine and human CD4^+^ T cells

Murine and human CD4^+^ T cells were labelled with CellTrace Violet (Life technologies) and cultured (200.000 cells/well) in either a 2D suspension (as a control) or the different hydrogels, as specified above. Human CD4^+^ T cells were cultured in RPMI10% medium (RPM1 1640 medium with Glutamax (Gibco) supplemented with 10% heat-inactivated FBS and 100 U/ml penicillin-streptomycin (Thermo Fisher Scientific)). Murine CD4^+^ T cells were cultured in RPMI10% supplemented with 1 mM sodium pyruvate (Gibco), 1X nonessential amino acids (Gibco), 10 mM HEPES (Gibco) and 50 μM 2-mercaptoethanol (Sigma-Aldrich). Functional grade monoclonal antibodies directed against mouse or human CD3 (anti-CD3, 1 µg/mL) and CD28 (anti-CD28, 1 µg/mL) (eBioscience) were incorporated into the matrices. After 5 days in culture at 37°C, CD4^+^ T cells were retrieved from the gels as specified above and stained for flow cytometry analysis. For intracellular staining of IFNγ, GolgiStop (containing Monensin, BD Biosciences) was added to the medium 4 hours before staining. In the case of NFC gels, GrowDase (2 µg/ml, UPM Biomedicals) was added simultaneously with GolgiStop.

### 4.6. Flow cytometry analysis of murine and human CD4^+^ T cells

Murine and human CD4^+^ T cells were stained with live/dead dye Zombie NIR (BioLegend) in PBS, followed by staining with fluorochrome-labelled antibodies in MACS buffer. Murine CD4^+^ T cells were stained with anti-CD4-APC (BioLegend) and anti-CD25-Pacific blue (BioLegend). Human CD4^+^ T cells were stained with anti-CD4-FITC (BioLegend) and anti-CD25-APC (BioLegend). For the assessment of human Treg cell differentiation, CD4^+^ T cells underwent intracellular staining using the Foxp3 / Transcription Factor Staining Buffer Set (eBioscience Thermofisher) with anti-FOXP3-PE (Biolegend). For both murine and human cells, intracellular IFNγ-staining was carried out using the Intracellular Fixation & Permeabilization Buffer Set (eBioscience Thermofisher) with anti-IFNγ-PECy7 (BD Biosciences for human and Biolegend for murine CD4^+^ T cells). Flow cytometry data were acquired using a BD LSRFortessa Cell Analyzer (BD Biosciences) with FACSDiva (BD Biosciences) software or a CytoFLEX Flow Cytometer (Beckman Coulter) with CytExpert software. Data were analyzed with FlowJo (Treestar, 10.6.2) or CytExpert software (2.4), respectively.

### 4.7. CD20 CAR-T cell generation and culture

The CD20 CAR construct (pBu-CD20–CAR) was generated by cloning single-chain variable fragments from anti-CD20 antibody rituximab into a pBullet vector containing a D8α−41BB-CD3-ζ signaling cassette. Phoenix-Ampho packaging cells were transfected with gag-pol (pHit60), env (P-COLT-GALV) and pBu-CD20–CAR using FugeneHD transfection reagent (Promega). Human peripheral blood mononuclear cells (PBMCs) were preactivated with 30 ng/mL anti-CD3 (OKT3, Miltenyi) and 50 IU/mL IL-2 (PeproTech) and subsequently transduced two times with viral supernatant in the presence of 6 μg/mL polybrene (Sigma) and 50 U/mL IL-2. Transduced T cells were expanded using 50 U/mL IL-2 and anti CD3/CD28 dynabeads (Thermo Fisher), and CD20–CAR-expressing cells were selected by treatment with 80 µg/mL neomycin. T cells were further expanded using rapid expansion protocol as described elsewhere [50].

### 4.8. CD20 CAR-T cell mediated cytotoxicity assays

Daudi cells (Burkitt lymphoma, DSMZ ACC 78) were cultured in RPMI 1640 GlutaMAX HEPES culture medium (Life Technologies) supplemented with 10% fetal bovine serum (FBS, Sigma) and 100 µg/mL penicillin–streptomycin (Life Technologies). For cytotoxicity assays, Daudi cells were labelled with CellTrace Violet dye (Invitrogen) and mixed with CD20 CAR-T cells at the effector to target (E:T) ratios indicated (50.000 Daudi cells/well). Cells were spun down and resuspended in the different hydrogels as specified above. After 24h, gels were digested and assessment of cell viability was performed by staining with 20 nM TO-PRO-3 (Thermo Scientific). The number of alive cells recovered from culture was quantified by flow cytometry, using Flow-count fluorospheres (Beckman Coulter).

### 4.9. Analysis of CAR-T cell proliferation and phenotype

CD20 CAR-T cells were labelled with CellTrace Violet (Life technologies) and cultured (50.000 cells/well) in either 2D or in hydrogels, as specified above. CAR-T cells were cultured in huRPMI2.5% medium (RPM1 1640 medium with GlutaMAX, supplemented with 2.5% pooled human serum, 100 µg/mL penicillin–streptomycin, 50 μM 2-mercaptoethanol and 50 IU/mL IL-2). Cells were stimulated with coated anti-CD3 (1.6 μg/ml) and soluble anti-CD28 (1 μg/ml) functional grade monoclonal antibodies (Miltenyi). After 4 days in culture at 37°C, CAR-T cells were recovered from the gels as described above. Cells were stained with anti-CD4 PECy7 (Invitrogen), anti-CD25 AF488 (Biolegend), and 20 nM TO-PRO-3 to assess viability. Samples were acquired in a LSRFortessa (BD Biosciences) with FACSDiva software and analyzed with FlowJo (Treestar, 10.6.2).

### 4.10. Statistical analysis

Data were analyzed using GraphPad Prism 10. Mean values with corresponding standard deviations (SD) are presented, and a minimum of three experiments were conducted for each group (refer to Figure Legends for additional details). Statistical significance was determined through ordinary one-way ANOVA with Tukey’s multiple comparison test unless otherwise stated. Significance levels are denoted by asterisks: *P ≤* 0.05 (*), *P ≤* 0.01 (**), or *P ≤* 0.001 (***) or *P* < 0.0001 (****).

## Acknowledgements

We thank Ralph G. Tieland for assistance with CAR-T cell cultures. We would like to thank Suze Jansen, Angelo Meringa and all members of the Coffer, Peperzak, Levato, Kranenburg and Lindemans group for helpful discussions. We thank the FACS facilities of the University Medical Center Utrecht (UMCU) and the Hubrecht Institute for their support. We would like to express our gratitude to all employees of NFCL Utrecht for their assistance with mice experiments. Funding: This work was supported in part by a Worldwide Cancer Research grant (Reference: 19-0371) and Marie S. Curie Co-fund RESCUE grant (Reference: 801540). The funding agencies played no role in the design, reviewing, or writing of the manuscript.

## Author Contributions

**Sonia Aristin Revilla:** Methodology, Formal analysis, Investigation, Writing - Original Draft, Review and Editing, Visualization. **Alessandro Cutilli:** Methodology, Formal analysis, Investigation. **Dedeke Rockx-Brouwer:** Investigation. **Cynthia Lisanne Frederiks:** Investigation. **Marc Falandt:** Methodology, Formal analysis, Investigation, Visualization. **Riccardo Levato:** Writing - Review and Editing, Supervision. **Onno Kranenburg:** Writing - Review and Editing, Supervision. **Caroline A. Lindemans:** Supervision. **Paul James Coffer:** Writing - Review and Editing, Supervision, Funding acquisition. **Victor Peperzak:** Writing - Review and Editing, Supervision. **Enric Mocholi:** Conceptualization, Methodology, Writing - Review and Editing, Supervision and Project Administration. **Marta Cuenca:** Conceptualization, Methodology, Formal analysis, Investigation, Writing - Original Draft, Review and Editing, Visualization, Supervision and Project Administration.

## Data availability

Data will be made available on request.

## List of abbreviations

ATMP: Advanced Therapy Medicinal Products
BME: basement membrane extract
CAR-T: chimeric antigen receptor T cell
CTLA-4: Cytotoxic T-Lymphocyte Antigen 4
ECM: extracellular matrix
IFNγ: interferon gamma
IGF-1: insulin growth factor 1
MFI: mean fluorescence intensity
NFC: nanofibrillar cellulose
PD-1: programmed cell death protein 1
TGF-β: transforming growth factor beta
TILs: tumor-infiltrating lymphocytes
TME: tumor microenvironment
Treg: regulatory T cell
VEGF: vascular endothelial growth factor.

