## Supplemental Figures for "Basement membrane hydrogels dampen CAR-T cell activation: nanofibrillar cellulose gels as alternative to preserve T cell function in 3D cell cultures"

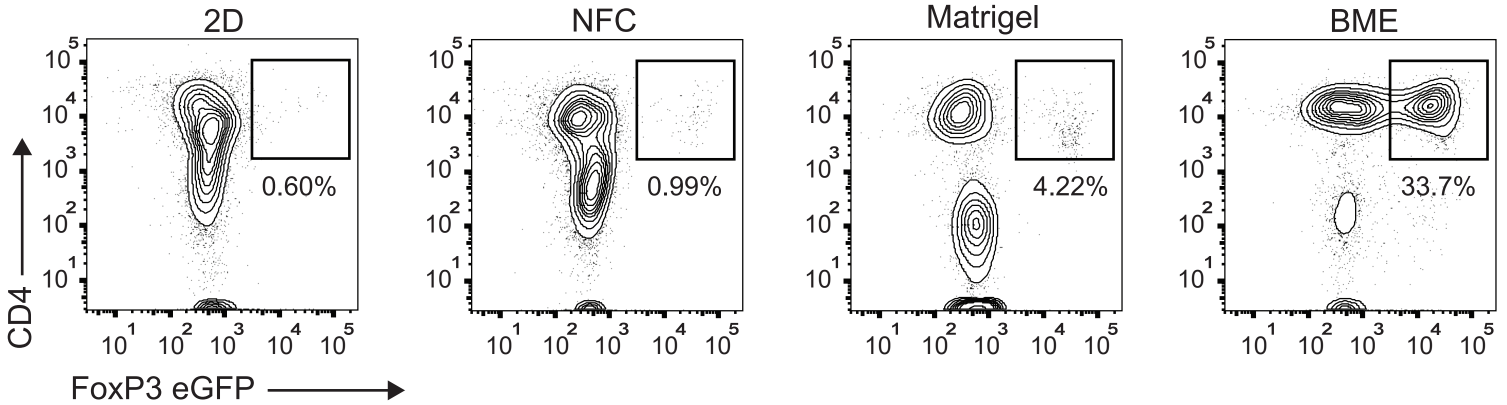


Supplemental Figure 1

Aristin Revilla S. *et al.*

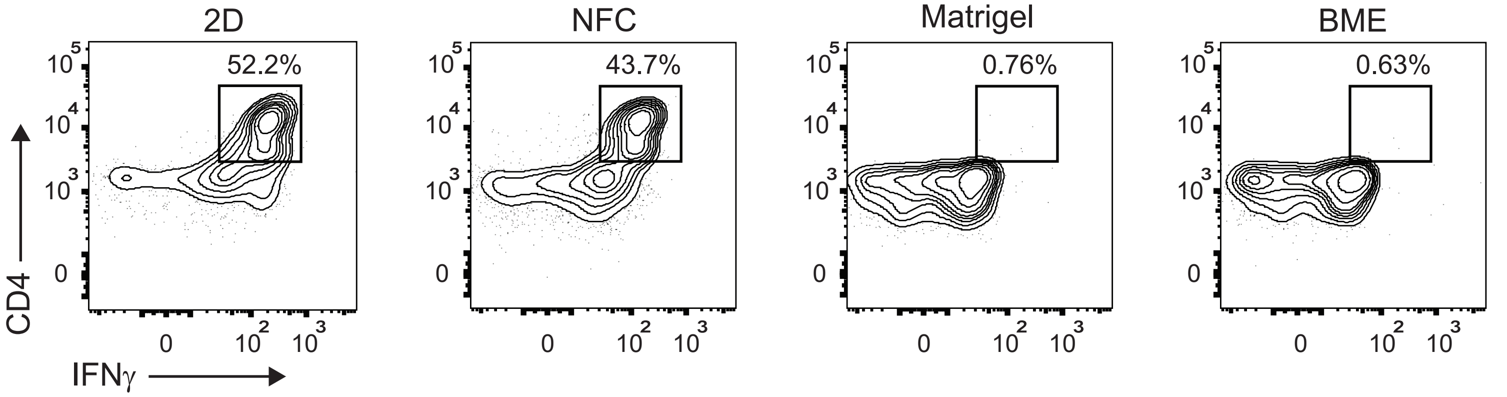
2D
NFC
Matrigel
BME
52.2%
43.7%
0.76%
0.63%
CD4
IFNγ

**Supporting Information**

**Supplemental Figure 1.** Increased Treg cell percentage when $CD4^+$ T cells are cultured in BME or Matrigel.

$CD4^+$ T cells were isolated from Foxp3eGFP mice and stimulated *ex-vivo* with anti-CD3 (1 μg/ml) and anti-CD28 (1 μg/ml) in 2D suspension (control) or embedded in different hydrogels (NFC, Matrigel and BME) for 5 days. Representative flow cytometry plots show the proportion of FoxP3 $eGFP^+CD4^+$ cells in each condition (gated on Alive cells). NFC, nanofibrillar cellulose; MG, Matrigel; BME, basement membrane extract.

**Supplemental Figure 2.** Matrigel and BME hinder $CD4^+$ T cell activation.

$CD4^+$ T cells were isolated from cord blood mononuclear cells and stimulated *ex-vivo* with anti-CD3 (1 μg/ml) and anti-CD28 (1 μg/ml) in 2D suspension (control) or embedded in different hydrogels (NFC, Matrigel and BME) for 5 days. Representative flow cytometry plots show the proportion of $IFNg^+CD4^+$ cells in each condition (gated on Alive cells). NFC, nanofibrillar cellulose; MG, Matrigel; BME, basement membrane extract.
